# Comparison of domain adaptation techniques for white matter hyperintensity segmentation in brain MR images

**DOI:** 10.1101/2021.03.12.435171

**Authors:** Vaanathi Sundaresan, Giovanna Zamboni, Nicola K. Dinsdale, Peter M. Rothwell, Ludovica Griffanti, Mark Jenkinson

## Abstract

Robust automated segmentation of white matter hyperintensities (WMHs) in different datasets (domains) is highly challenging due to differences in acquisition (scanner, sequence), population (WMH amount and location) and limited availability of manual segmentations to train supervised algorithms. In this work we explore various domain adaptation techniques such as transfer learning and domain adversarial learning methods, including domain adversarial neural networks and domain unlearning, to improve the generalisability of our recently proposed triplanar ensemble network, which is our baseline model. We evaluated the domain adaptation techniques on source and target domains consisting of 5 different datasets with variations in intensity profile, lesion characteristics and acquired using different scanners. For transfer learning, we also studied various training options such as minimal number of unfrozen layers and subjects required for finetuning in the target domain. On comparing the performance of different techniques on the target dataset, unsupervised domain adversarial training of neural network gave the best performance, making the technique promising for robust WMH segmentation.

## 1. Introduction

White matter hyperintensities of presumed vascular origin (WMHs, also known as white matter lesions) are bright localised regions on T2-weighted and FLAIR images. They are commonly found in elderly subjects, however, they have also been related to various neurodegenerative (e.g. dementia, including Alzheimer’s disease) and cerebrovascular diseases (e.g. stroke) (Wardlaw et al., 2013). Automated WMH segmentation is essential for further understanding the clinical impact of WMHs in a large population. Various methods using hand-crafted features have been used for WMH segmentation (Caligiuri et al., 2015), and in recent years, deep learning (DL) models are being increasingly used and have been shown to outperform traditional methods (Rachmadi et al., 2018; Kuijf et al., 2019). Many of the existing methods (using either hand-crafted features or DL models) were trained with a large amount of manual labels (Wang et al., 2012; Admiraal-Behloul et al., 2005; Ghafoorian et al., 2016) and/or evaluated on specific population group (Wang et al., 2012; Gibson et al., 2010; De Boer et al., 2009; Steenwijk et al., 2013; Jeon et al., 2011; Hong et al., 2020; Park et al., 2018), acquired with the same scanner/protocol or validated on isotropic or axial acquisition images (Ghafoorian et al., 2017a; Kuijf et al., 2019). However, in the real-world scenario, most of the clinical datasets are small in size, acquired using various protocols and scanners, and from people with diverse demographic and pathological characteristics. In addition, specially in these datasets, limited amount or non-availability of manual segmentations constrains the training and segmentation performance of the model, especially due to the problem of overfitting. It is therefore very challenging to achieve robust performance metrics for segmentation of WMHs across datasets in the presence of such variations in image characteristics, lesion load, and availability of training data.

Several methods have been proposed for making models more adaptable to various ‘*domains*’ (e.g. different scanners or acquisition protocols). These include reducing the variance in the image-level characteristics (Bordin et al., 2020) (induced by the scanner and acquisition protocol), estimating site effects to correct the measurements derived from the images (Fortin et al., 2018), by improving model generalisability (Ganin et al., 2016; Tzeng et al., 2015) (so that it is not affected by differences in intensity distributions or spatial resolution), or a combination of the above. Commonly used techniques to improve model generalisability include data augmentation (Shorten and Khoshgoftaar, 2019), and the use of ensemble networks (with different initialisations (Li et al., 2018) or planes (Prasoon et al., 2013)), which have been shown to be resistant to over-fitting (Krizhevsky et al., 2012; Simonyan and Zisserman, 2014; Kamnitsas et al., 2017; Winzeck et al., 2019), which can occur with more complex models (Opitz and Maclin, 1999). However, these techniques cope mostly with minor variance in dataset characteristics within a domain and hence might not be sufficient for generalising across datasets obtained from different sources/domains.

Domain adaptation (DA) methods address the issue of discrepancies in the data distributions obtained from various domains that affect the robust performance of the model (Ben-David et al., 2010). DA methods, in general, aim to transfer the knowledge from a *source* domain to a *target* domain by leveraging the invariant features across different domains (Wilson and Cook, 2019; Pan and Yang, 2009). Various DA techniques used so far include minimising a distance metric of domain variance (Long et al., 2013; Pan et al., 2010; Wang and Schneider, 2014), using transferable features for creating intermediate feature representation between domains (Yosinski et al., 2014) and transfer learning (Pan and Yang, 2009; Yosinski et al., 2014). Within DA frameworks, the restriction posed by limited availability of manually labelled data for training has been addressed by proposing various semi-supervised (Cheng and Pan, 2014; Yao et al., 2015; Saito et al., 2019), self-labelling (Saito et al., 2017; Zou et al., 2019) and pseudo-labelling (Inoue et al., 2018) methods. Techniques such as self- and pseudo-labelling use small amount of labelled data along with large amount of unlabelled data to improve model performance (Lee et al., 2013). Hence, given the wide variations in lesion characteristics, contrast variations (e.g. GM voxels vs WMHs) and location priors (e.g. normal ventricle lining vs WMHs), these techniques could bias WMH segmentation results, especially in small non-representative datasets.

In *transfer learning* (TL) (Pan and Yang, 2009), one of the commonly used supervised DA techniques, the initial convolutional layers (domain invariant low-level features) are generally kept constant or *frozen*, while the final layers (task/domain specific high-level features) are fine-tuned on the target datasets. Fine-tuning of the pre-trained models has been shown to improve the performance on the target datasets (Tajbakhsh et al., 2016), especially when the intensity characteristics of images between domains are more similar (Yosinski et al., 2014; Wilson and Cook, 2019). TL has been applied for medical image segmentation tasks (Tajbakhsh et al., 2016), including lesion segmentation (Ghafoorian et al., 2017b). However, TL is limited by the fact that the training on different domains occurs separately and hence cannot combine features from both domains while training. Also, in addition to determining the layers to fine-tune, another crucial consideration often encountered while fine-tuning is the number of target training subjects required. This is because the performance of TL has been shown to rely (although to a less extent than training the model from scratch) on the amount of labelled target training data (Tajbakhsh et al., 2016).

Another successful DA approach, *unsupervised domain adversarial training* (Ganin et al., 2016; Tzeng et al., 2015; Lee et al., 2019; Fernando et al., 2013; Kouw and Loog, 2019; Wang and Deng, 2018) relies on domain invariant features to achieve good domain adaptation. Several adversarial training methods have been proposed, including the recent ones based on discriminator framework (Tzeng et al., 2017), partial transfer learning (Cao et al., 2018) (assuming that the target domain dataset is a subset of the source domain) and using associations between the source and target domains (e.g. increasing correlation/covariance, subspace alignment) (Haeusser et al., 2017; Fernando et al., 2013; Long et al., 2017a; Sun and Saenko, 2016). One of the earlier and commonly used adversarial training approaches, domain adversarial training of neural network (DANN) (Ganin and Lempitsky, 2015; Ganin et al., 2016) has been applied to several baseline architectures (Ganin et al., 2016; Schoenauer-Sebag et al., 2019) and is explored on multiple datasets (Gallego et al., 2020). The DANN technique has also been used in various comparative analyses (Tzeng et al., 2015, 2017; Zellinger et al., 2017) and has been proven to be one of the successful models for task-specific domain adaptation (Zellinger et al., 2017; Long et al., 2017b). The DANN model consists of a feature extractor network with a domain predictor and a label predictor. The adversarial training of the domain predictor is achieved using a *gradient-reversal layer*, placed between the feature extractor and the domain predictor, optimising the features and shared weights. This layer maximises the domain prediction loss, thus minimising the shift between the domains, while simultaneously making the model discriminative towards the main task of segmentation label prediction. While DANN was originally proposed in an unsupervised manner with respect to target domain (i.e. segmentation labels required only for source domain), it has also been shown to be beneficial under semi-supervised setting (i.e. using a fraction of target domain segmentation labels) (Ganin et al., 2016).

An alternative domain unlearning (DU) approach was proposed for domain and task adaptation using an iterative framework (Tzeng et al., 2015) (and recently adapted for unlearning scanner-related information between domains in (Dinsdale et al., 2020)). The method involved learning the domain prediction for a fixed feature representation and then minimising the domain shift between features resulting in a maximal domain confusion that is equally uninformative across domains.

In this work, we explore various domain adaptation techniques such as TL and adversarial adaptation methods including DANN and DU for obtaining a good WMH segmentation across various datasets, and to perform well irrespective of differences in data characteristics. We used a triplanar ensemble network (TrUE-Net), proposed in our recent work (Sundaresan et al., 2020) as a baseline model. Our objective is to adapt our baseline model to a different domain consisting of small dataset(s). In addition, when applying TL, we addressed two main issues while fine-tuning a model on a target dataset with limited training subjects: determining (1) the optimal layer of the model to start fine-tuning and (2) the minimal number of training subjects for reliable segmentation. We performed our experiments on 5 different datasets (including 3 publicly available datasets) with different acquisition and lesion characteristics, grouped into source and target domains. We experimented with several test cases involving different DA techniques on the source-trained model and training the model directly on the target domain, to comprehensively study both innate and adapted performances of the model for WMH segmentation.

## 2. Materials and methods

### 2.1. Datasets used

#### Neurodegenerative cohort (NDGEN)

The dataset, used in Zamboni et al. (2013), includes MRI data from 9 subjects with probable Alzheimer’s Disease, 5 with amnestic mild cognitive impairment and 7 cognitively healthy control subjects (age range 63 - 86 years; mean age 77.1 ± 5.8 years; median age 77 years; F:M = 10:11). Total brain volume range: 1189282 - 1614799 mm^3^, median: 1424669 mm^3^. Manual segmentation was available for all datasets (WMH load range: 1878 - 89259 mm^3^, median: 20772 mm^3^). The images were acquired using a 3T Siemens Trio Scanner, with FLAIR (TR/TE = 9000/89 ms, flip angle 150°, FOV 220 mm, voxel size 1.1 × 0.9 × 3 mm, matrix size 256 × 256 × 35 voxels) and T1-weighted acquisitions (3D MP-RAGE sequence, TR/TE = 2040/4.7 ms, flip angle 8°, FOV 192 mm, voxel size 1 mm isotropic, matrix size 174 × 192 × 192 voxels).

#### Vascular cohort - Oxford Vascular Study (OXVASC)

The dataset consists of 18 participants in the OXVASC study (Rothwell et al., 2004), who had recently experienced a minor non-disabling stroke or transient ischemic attack (age range 50 - 91 years; mean age 73.27 ± 12.32 years; median age 75.5 years; F:M = 7:11). Total brain volume range: 1290926 - 1918604 mm^3^, median: 1568233 mm^3^. Manual segmentation was available for all datasets (WMH load range: 3530 - 83391 mm^3^, median: 16906 mm^3^). The images were acquired using a 3T Siemens Trio Scanner, with FLAIR (TR/TE = 9000/88 ms, flip angle 150°, voxel size 1 × 3 × 1 mm, matrix size 174 × 52 × 192 voxels) and T1-weighted acquisitions (3D MP-RAGE sequence, TR/TE = 2000/1.94 ms, flip angle 8°, voxel size 1 mm isotropic, matrix size 208 × 256 × 256 voxels).

#### MICCAI WMH Segmentation Challenge training Dataset (MWSC)

The dataset consists of 60 subjects from three different sources (20 subjects each) provided as training sets for the challenge (Kuijf et al., 2019) (http://wmh.isi.uu.nl/): UMC Utrecht, NUHS Singapore and VU Amsterdam. The brain volume ranges: 1257820 - 1844920 mm^3^(median – 1473389 mm^3^) for UMC Utrecht, 1147248 - 1532268 mm^3^(median: 1351325 mm^3^) for NUHS Singapore and 1219614 - 1787321 mm^3^(median: 1441201 mm^3^) for VU Amsterdam. Manual segmentations were available for all three datasets, with an additional exclusion label provided for other pathologies. We included these masks as parts of non-lesion tissue during both training and testing. The WMH volume ranges (excluding other pathologies) are 845 - 74991 mm^3^(median: 26240 mm^3^) for UMC Utrecht, 786 - 61332 mm^3^(median: 17795 mm^3^) for NUHS Singapore and 1522 - 43528 mm^3^(median: 6015 mm^3^) for VU Amsterdam. FLAIR and T1-weighted images were available for this dataset (for more details regarding MRI acquisition parameters, refer to http://wmh.isi.uu.nl/).

### 2.2. Data preprocessing

We reoriented FLAIR and T1-weighted images to the standard MNI space, performed skull-stripping with FSL BET (Smith, 2002) and bias field correction using FSL FAST (Zhang et al., 2001). We registered the T1-weighted image to the FLAIR using rigid-body registration using FSL FLIRT (Jenkinson and Smith, 2001) and cropped the field of view (FOV) close to the brain and applied Gaussian normalisation to the intensity values. For axial, sagittal and coronal slices, we resized the extracted slices to dimensions of 128 × 192, 192 × 120 and 128 × 80 voxels respectively, using bilinear interpolation.

### 2.3. Baseline method: Triplanar U-Net Ensemble Network (TrUE-Net) architecture

As a baseline model, we used the triplanar ensemble architecture^2^ proposed in Sundaresan et al. (2020). Briefly, as shown in figure 1, the TrUE-Net architecture consists of three 2D U-Nets, one for each plane, taking FLAIR and T1 slices as input channels. We trimmed the depth of the classic U-Net (Ronneberger et al., 2015) in each plane to a depth of 3-layers (figure 2a), given the small size of lesions. In the ensemble model, we trained the U-Nets in each plane independently using 2D slices extracted in each plane. We used a combination of weighted cross-entropy (refer to Sundaresan et al. (2020) for more details) and Dice loss functions in order to overcome the effect of class imbalance between WMHs and healthy tissue. During testing, the predictions were obtained as 2D softmax output score maps for slices in each plane and were later assembled into 3D volumes and resized to the original dimensions. We then averaged the 3D volumes to get the final probability volume for the triplanar architecture.

**Figure 1:**
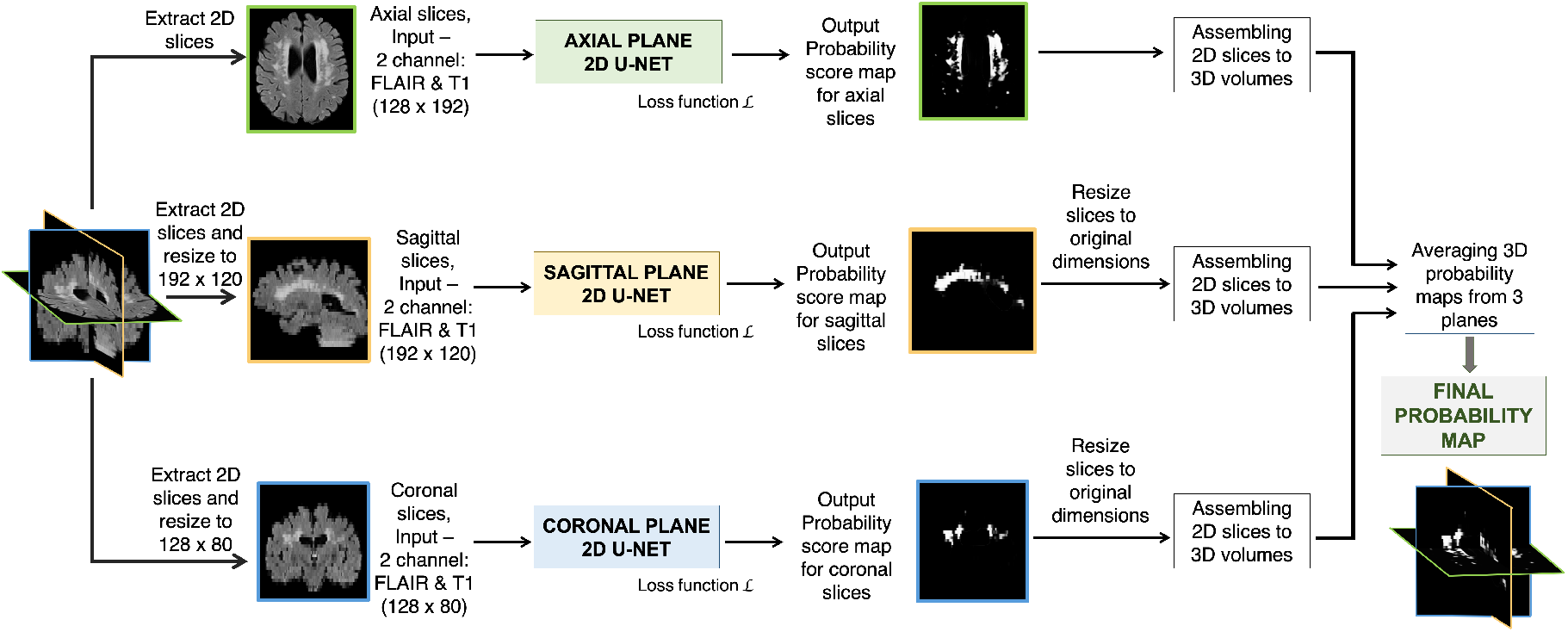
Baseline architecture: Triplanar U-Net ensemble network (TrUE-Net) Sundaresan et al. (2020).

**Figure 2:**
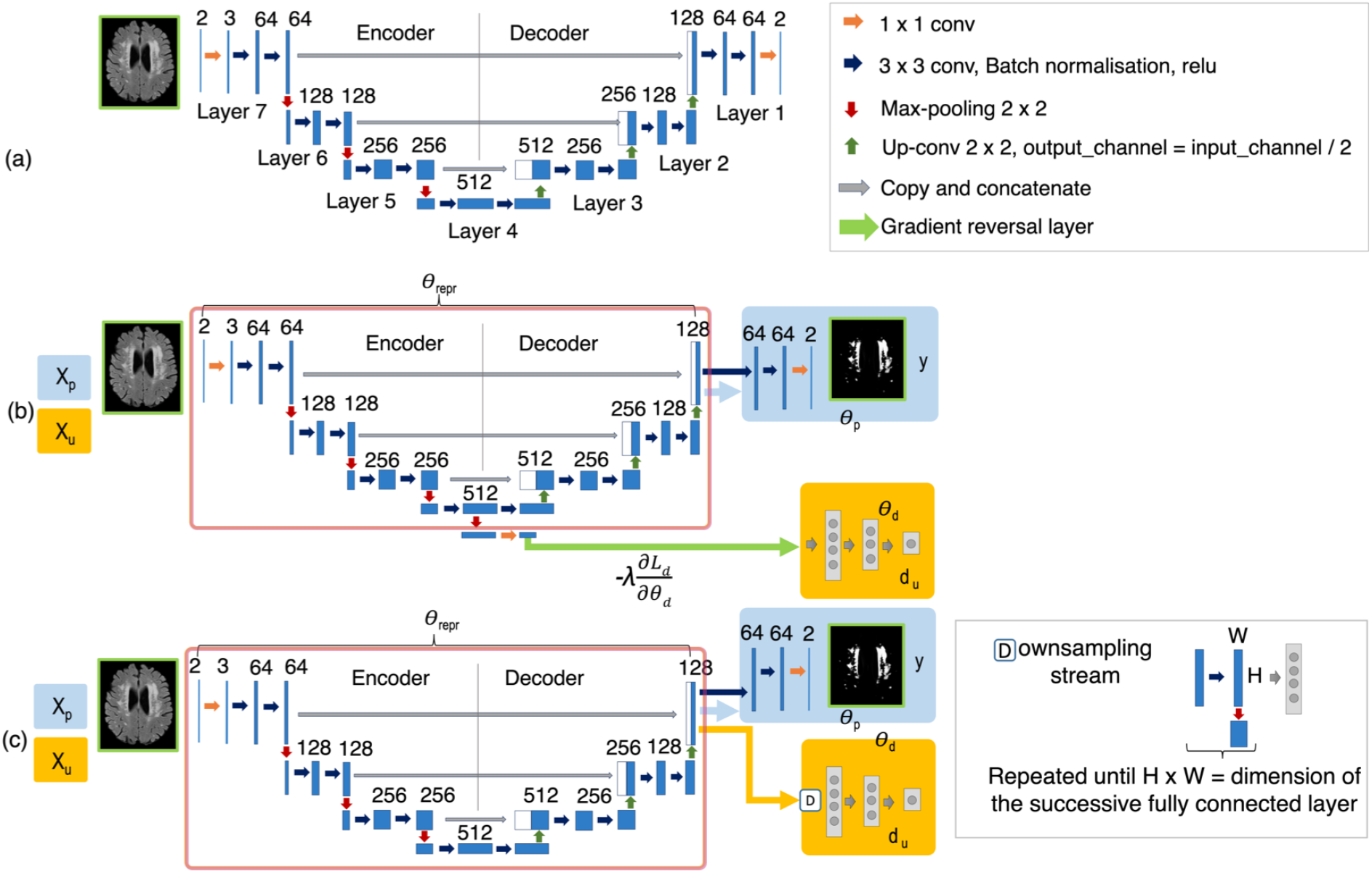
Transfer learning (TL) framework, domain adversarial neural network (DANN) and domain unlearning (DU) architectures. (a) Layer numbers indicated on the baseline model for the TL case (numbered from decoder end indicating the order of fine-tuning), (b) DANN and (c) DU architectures, illustrating feature extractor (red box), lesion label predictor (blue) and domain predictor (orange) with corresponding training parameters *θ*_*repr*_, *θ*_*p*_ and *θ*_*d*_. The models take input features *X*_*p*_ and input domain information *X*_*u*_ and predicts output labels y, while unlearning output domains *d*_*u*_. The DU model updates the label predictor, feature extractor and domain predictor in a sequential manner, while label prediction and domain unlearning occur simultaneously in DANN. For all the cases, only the axial U-Net is shown; note that sagittal and coronal models were modified in a similar manner.

### 2.4. Comparison of domain adaptation techniques

We studied the performance of various DA techniques using the following test cases on the target test dataset (refer to section 2.6) against the case where the model is trained directly on the target training dataset. Due to the non-Gaussian distribution of the performance metrics data in most cases (Shapiro-Wilks test), we performed non-parametric statistics using Wilcoxon signed rank test on the metrics described in section 2.7, between the individual pairs of test cases for comparison.

#### 2.4.1. Case 1: Train on the source domain and apply directly to the target domain

In order to determine the inherent generalisability of TrUE-Net, we trained the model on the source domain training datasets (training parameters in section 2.5) and tested the model directly on the target domain test datasets.

#### 2.4.2. Case 2: Transfer the model trained on the source domain to the target domain with fine-tuning

We trained TrUE-Net (training details in section 2.5) on the source domain datasets to get the source pre-trained model. We then fine-tuned the model by training it on the target dataset starting from the decoder end. For fine-tuning we used a smaller learning rate schedule (initially 1 × 10^−4^, reduced by a factor 1 × 10^−1^every 2 epochs, until it reaches 1 × 10^−6^). Figure 2a shows the layer numbers for the U-Net model. Given *L* layers in total, ‘fine-tuning *i* layers’ means that *L* − *i* layers before *i* were frozen and the layers from *i* towards the decoder end were fine-tuned. For each fine-tuning, we increased the number of target training subjects, from 2 to 18 in steps of 2, and measured the performance of each fine-tuned model. Finally, we determined the best starting point for fine-tuning the model, and the optimal number of training data to obtain the best performance on the target dataset. Since TL involves both the domains, in addition to the existing DA cases, we also compare the TL case with the case where the baseline model is trained on the source and target training datasets combined together (refer to section 4 in the supplementary material).

#### 2.4.3. Case 3: Unsupervised domain adversarial training (DANN) on the source and target domains

We implemented DANN (Ganin et al., 2016) as shown in figure 2b by adding a domain predictor to the baseline TrUE-Net model. We added the domain predictor to the coarsest level after maximum levels of pooling (with 512 channels) at the end of the encoder, since it has high-level features with domain-specific information. We added a 2 × 2 max-pooling layer and 1 × 1 projection layers to 128 and 64 channels before the domain predictor. The domain predictor consists of three fully connected (FC) layers (with 1024, 512 and 32 nodes) alternating with two dropout layers (*P*_*drop*_ = 0.2%), followed by a softmax layer. We added a gradient reversal layer between the feature extractor and the domain predictor, leading to adversarial training with respect to domain prediction. In this model, the domain predictor makes the model domain invariant by considering data from both domains, while the lesion label predictor optimises the model for accurate WMH segmentation. Hence, the domain predictor requires only domain labels for both source and target datasets, while the lesion label predictor requires manual segmentations from source datasets only, making the model unsupervised with respect to the target domain. We tested the DANN model on the test datasets from the source and target domains (table in figure 3), and measured model performance in each domain individually.

**Figure 3:**
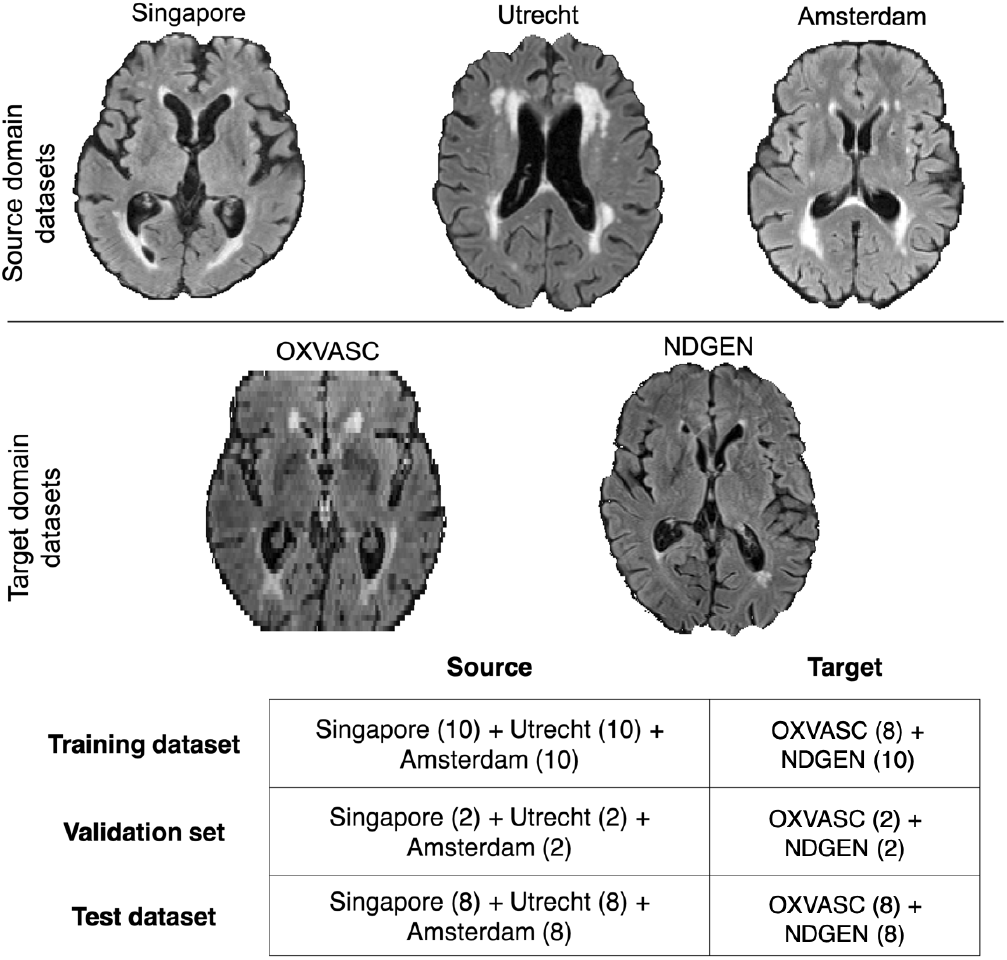
Sample axial slices shown from source (top row) and target (middle row) domains. The splits for training, validation and test datasets in source and target domains are provided in the table (bottom panel).

#### 2.4.4. Case 4: Semi-supervised domain adversarial training (semi-DANN) on source and target domains

We trained the DANN model (figure 2b) in a semi-supervised manner, wherein manual segmentations from a fraction of target training data were used in addition to source training datasets for training the lesion label predictor. The remaining target training data is used only for domain prediction. One of the main advantages of the unsupervised DANN model is that it does not require manually labelled data from the target dataset. Even then, our main aim for exploring the semi-DANN, despite the additional manual labelling effort, was to observe if there was any significant improvement over unsupervised DANN. Hence, we used a minimal proportion (25%, chosen empirically, which amounts to 4 subjects) of the labelled target training data, in addition to the source training dataset, for training the lesion label predictor. We tested the model on the source and target domain test datasets individually.

#### 2.4.5. Case 5: Iterative domain unlearning (DU) to remove scanner-bias between source and target domains

The DU model^3^(Dinsdale et al., 2020) is based on the iterative unlearning framework (Tzeng et al., 2015) for adversarial adaptation. The model, rather than using a gradient reversal layer, optimises two opposing loss functions in three sequential steps: (1) updating the feature representation and the lesion label predictor, (2) maximising the performance of a domain predictor given the fixed feature representation, and (3) updating the feature representation in order to maximally confuse the domain classifier. As in the case of the unsupervised DANN (case 3), only domain labels are required for the target dataset, while the lesion label predictor uses manual segmentations of the source dataset only. As shown in figure 2c, we consider the final two 3 × 3 convolutional layers and the final softmax layer as our label predictor. The domain predictor, placed after the final decoder layer of the U-Net, consists of repetitive 2 × 2 max-pooling layers until the last layer dimensions match the first FC layer, followed by three FC layers (with 468, 96 and 32 nodes) alternating with two dropout layers (*P*_*drop*_ = 0.2%), followed by a softmax layer. Note that while we added the domain predictor at the end of the encoder in DANN, we added the domain predictor at the end of decoder (the same point as label predictor) in DU (figure 2). This is because, in DANN, the domain unlearning happens simultaneously with shared weights between the predictors and hence we focus on the layer with coarsest features (showing overall WMH distribution) that is more domain specific. On the other hand, in DU, the training happens sequentially for each predictor (while freezing the other predictor) and hence we add the domain predictor directly at the point where the generalisability is most desirable for WMH segmentation.

#### 2.4.6. Train on the target domain from scratch and apply to the target domain

For this case, we trained the TrUE-Net model on the target training dataset and tested the model on the target test dataset. This case is expected to perform better than other DA cases on the target test dataset (for the given training options and data) since it is not required to cope with domain variance. Hence, this case is included for comparison purposes, though it does not improve model generalisability.

### 2.5. Implementation details

We implemented the networks in Python 3.6 using Pytorch 1.2.0. The baseline network (TrUE-Net), for source-trained, target-trained and TL (pretraining) cases, was trained on an NVIDIA Tesla V100, taking 5 mins (for 3 planes) per epoch for ≈ 15,000 samples with the training/validation split of 90/10%. We used the Adam Optimiser with *ϵ*=10^−4^. We empirically chose a batch size of 8, and an initial learning rate of 1×10^−3^and reducing it by a factor 1 × 10^−1^ every 2 epochs, until it reaches 1 × 10^−5^, after which we maintain a fixed learning rate value (for more details, refer to Sundaresan et al. (2020)). Data augmentation was applied in an online manner using translation (x/y-offset ∈ [−10, 10]), rotation (*θ* ∈ [−10, 10]), random noise injection (Gaussian, *μ* = 0, *σ*^2^ ∈ [0.01, 0.09]) and Gaussian filtering (*σ* ∈ [0.1, 0.3]), increasing the dataset by a factor of 10 and 6 for axial and sagittal/coronal planes respectively. The hyperparameter values for the data augmentation transformations were randomly sampled from the closed intervals specified above using a uniform distribution. Additionally, for the domain predictor in DANN/semi-DANN and DU, we trained with the Momentum optimiser (momentum value of 0.9) and Adam optimiser respectively. We used a batch size of 8, with 50 epochs for pretraining and a criterion based on a patience value (number of epochs to wait for progress on validation set) of 25 epochs to determine model convergence for early stopping (converged at around 90 epochs for all cases). We used the learning rates of 10^−3^and 10^−4^for DANN/semi-DANN and DU respectively. These training hyper parameters were chosen empirically. In DU, we used a *β* value of 50 (a factor used for weighting the domain confusion (Dinsdale et al., 2020)). For determining the *β* value, we experimented with different values starting from 20 to 60 in steps of 10 and chose the value of 50, since it provided a domain accuracy value closer to 50% (indicating maximal confusion of domains) on the validation dataset. The DANN/semi-DANN and DU networks were trained on an NVIDIA Tesla V100, taking ≈ 10 and 7 mins per epoch respectively with the training/validation split of 90/10%.

### 2.6. Source and target domain datasets

The datasets used in this work were acquired using different scanners and sequences and therefore have different intensity characteristics and resolutions. For performing our domain adaptation (DA) experiments, we classified the available datasets into two domains. Rather than considering each dataset as an individual domain, we considered only two domains (source and target) for our experiments. This is because the datasets have varying degrees of similarity among them and also, given the limited number of subjects for each dataset (for training and testing), treating them as individual domains would be difficult and give unreliable results. For deciding the source and target datasets, we determined the homogeneity of image-level characteristics among the above 5 datasets, using a domain discriminator network. To this aim, we trained a domain discriminator model on the above datasets and determined the domain misclassifications among these datasets using a confusion matrix (for more details on this experiment and results, refer to section 1 of supplementary material). Based on the results, we considered the MWSC dataset (3 cohorts) as our *source* domain datasets, and the combination of OXVASC and NDGEN as our *target* domain datasets. Examples of the source and target domain datasets, along with the training/validation/test data split for above test cases is shown in figure 3.

### 2.7. Performance evaluation metrics

We used the following performance metrics:

- Dice Similarity Index (SI) = 2 × (true positive WMH voxels) / (true WMH voxels + positive WMH voxels).
- Voxel-wise true positive rate (TPR), the ratio of the number of true positiveWMHvoxels to the number of true WMH voxels.
- Voxel-wise false positive rate (FPR), the number of false positive WMH voxels divided by the number of non-WMH voxels.
- Cluster-wise TPR, the number of true positive WMH clusters (determined using 26-connected neighbourhood) divided by the total number of true WMH clusters.
- Absolute volume difference (AVD) between manual and predicted segmentations in percentage
- Cluster-wise F1-measure = 2 × (cluster-wise TPR × cluster-wise precision)/ (cluster-wise TPR + cluster-wise precision), where cluster-wise precision is the number of true positive WMH clusters divided by the total number of detected WMH clusters.

For each metric we then performed Wilcoxon signed rank test between the individual pairs of test cases.

## 3. Results

Figure 4 shows results for all the test cases of the domain adaptation experiments for a sample high lesion load test subject from the OXVASC dataset (results on a low lesion load test subject are shown in figure S3 in the supplementary material). Table 1 reports the performance values of the test cases, while figure 5 shows the boxplots of the performance metrics. For the results of Wilcoxon signed rank tests comparing the evaluation metrics for each pair of test cases, refer to table 2. For the DANN, semi-DANN and DU models, figure 7 shows the visualisation of the feature representations using t-SNE plots (Van der Maaten and Hinton, 2008) at the layer before the label predictor with and without domain adaptation.

**Figure 4:**
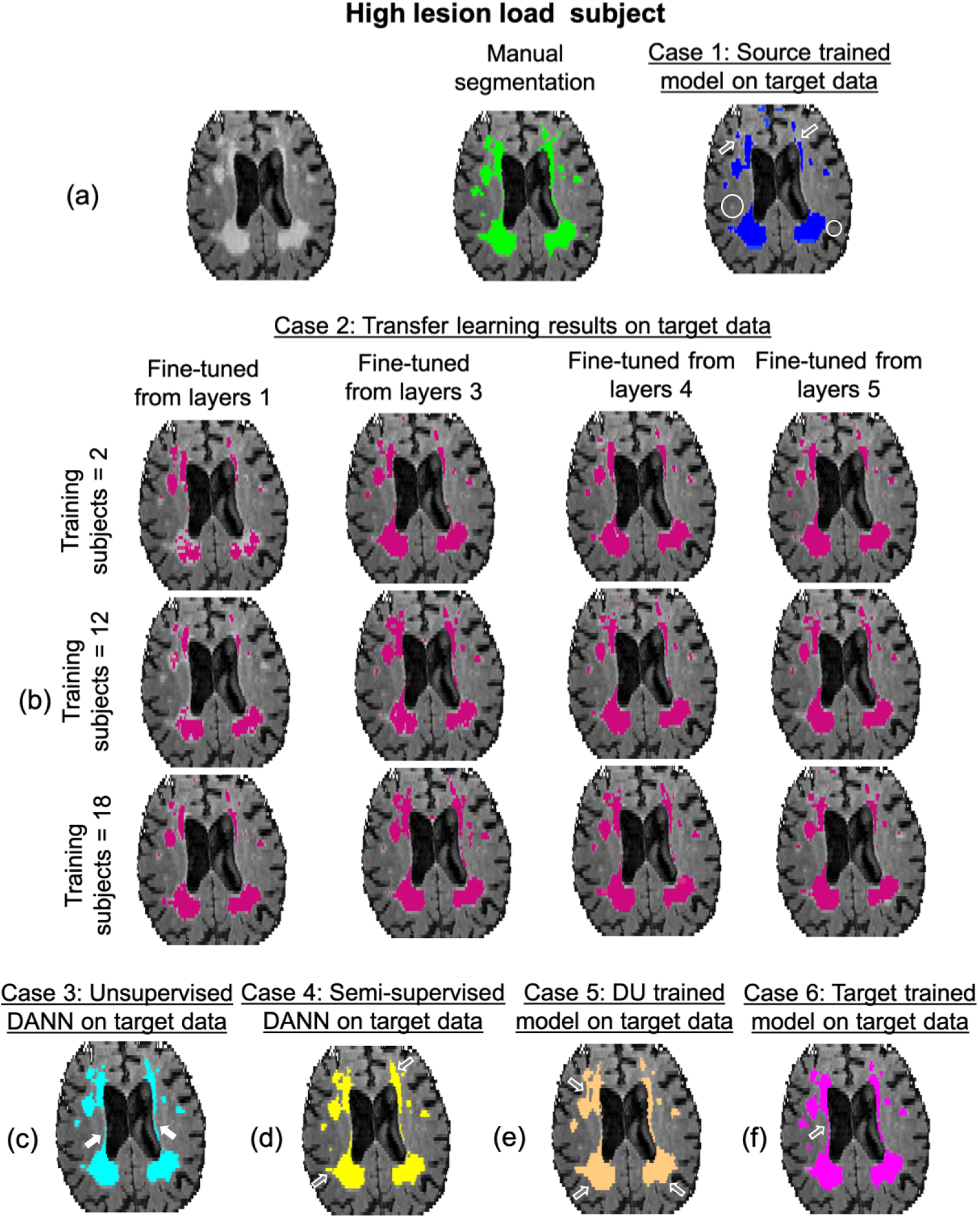
Sample results of domain adaption experiment test cases: (a) Source-trained model, (b) TL models, (c) unsupervised DANN, (d) semi-supervised DANN (semi-DANN), (e) DU and (f) target-trained model on a high lesion load subject from the OXVASC dataset (target domain), along with the manual segmentation. The over/under-segmented regions of periventricular WMHs are indicated by hollow arrows, the correctly predicted regions by filled arrows and missed deep WMHs are shown in circles.

**Table 1:** Comparison of performance metrics for 5 DA test cases against the target-trained case on the target test dataset (from NDGEN and OXVASC) (median and interquartile range (IQR) values provided; the best median value(s) for each performance metric is highlighted in bold).

| Perf. metrics | Case 1 | Case 2 | Case 3 | Case 4 | Case 5 | Target-trained |
| --- | --- | --- | --- | --- | --- | --- |
| SI value ↑ | 0.80<br>(0.68-0.86) | **0.89**<br>**(0.63-0.97)** | 0.78<br>(0.66-0.92) | 0.84<br>(0.76-0.92) | 0.77<br>(0.70-0.83) | 0.88<br>(0.84-0.98) |
| AVD (%) ↓ | 51.8<br>(32.7-71.9) | 40.3<br>(12.4-54.5) | 17.1<br>(7.7-35.2) | **12.8**<br>**(6.2-22.1)** | 19.7<br>(13.9-29.3) | 5.1<br>(2.5-14.1) |
| Cluster-wise F1-measure ↑ | 0.56<br>(0.43-0.63) | 0.62<br>(0.58-0.66) | 0.65<br>(0.55-0.81) | **0.79**<br>**(0.71-0.83)** | 0.69<br>(0.58-0.75) | 0.82<br>(0.58-0.98) |
| Cluster-wise TPR ↑ | 0.71<br>(0.59-0.85) | 0.83<br>(0.72-0.87) | **0.86**<br>**(0.71-0.98)** | 0.81<br>(0.77-0.90) | 0.74<br>(0.64-0.83) | 0.89<br>(0.66-0.99) |
| Voxel-wise TPR ↑ | 0.77<br>(0.70-0.89) | 0.84<br>(0.71-0.87) | **0.87**<br>**(0.82-0.94)** | **0.87**<br>**(0.78-0.91)** | 0.85<br>(0.67-0.98) | 0.97<br>(0.82-0.98) |
| Voxel-wise FPR ↓ | 1.8<br>(1.3-5.2) $\times 10^{-4}$ | 1.6<br>(0.9-3.4) $\times 10^{-4}$ | **0.8**<br>**(0.3-1.6) $\times 10^{-4}$** | 1.3<br>(0.5-3.3) $\times 10^{-4}$ | 1.7<br>(1.0-2.8) $\times 10^{-4}$ | 1.2<br>(0.6 - 4.7) $\times 10^{-4}$ |

**Figure 5:**
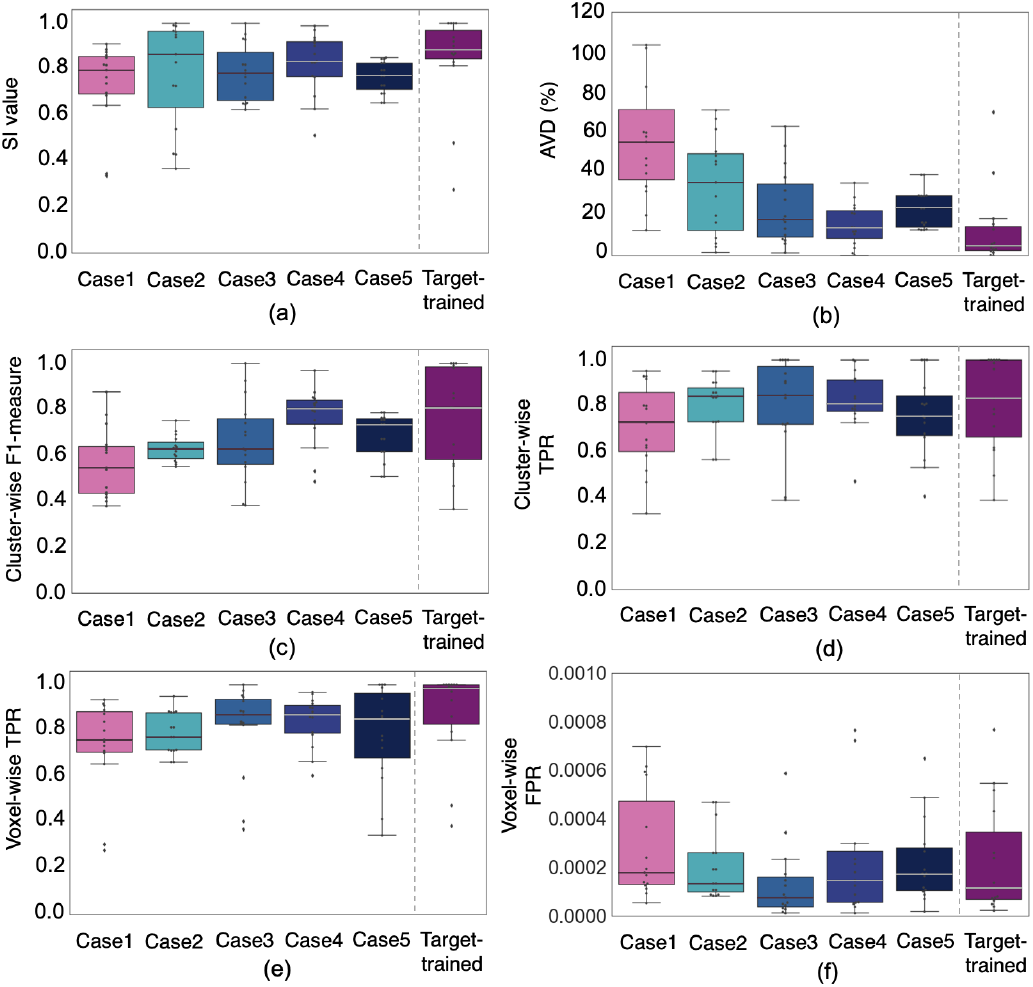
Boxplots of performance metrics obtained for the 5 test cases of the domain adaptation experiment, shown against the target-trained case, on the target test dataset (OXVASC + NDGEN) - (a) SI values, (b) AVD (%), (c) cluster-wise F1-measure, (d) cluster-wise TPR, (e) voxel-wise TPR and (f) voxel-wise FPR values.

**Table 2:** Results of Wilcoxon signed rank test results between individual combinations of test cases on the target dataset (p *<* 0.05 highlighted in bold). Cl. F1 - Cluster-wise F1 measure, Cl. TPR - Cluster-wise true positive rate (TPR), Vox. TPR - Voxel-wise TPR, Vox. FPR - Voxel-wise false positive rate.

|  | Case 2 | Case 3 | Case 4 | Case 5 | Target-trained |
| --- | --- | --- | --- | --- | --- |
| Case 1 | SI: **p = 0.04**<br>AVD: p = 0.06<br>Cl. F1: p = 0.18<br>Cl. TPR: p = 0.12<br>Vox. TPR: p = 0.98<br>Vox. FPR: **p = 0.04** | SI: p = 0.18<br>AVD: **p = 0.002**<br>Cl. F1: **p = 0.03**<br>Cl. TPR: **p = 0.004**<br>Vox. TPR: **p = 0.009**<br>Vox. FPR: **p < 0.001** | SI: **p = 0.002**<br>AVD: **p < 0.001**<br>Cl. F1: **p < 0.001**<br>Cl. TPR: **p = 0.003**<br>Vox. TPR: **p = 0.03**<br>Vox. FPR: p = 0.68 | SI: p = 0.64<br>AVD: **p < 0.001**<br>Cl. F1: **p = 0.02**<br>Cl. TPR: p = 0.05<br>Vox. TPR: p = 0.21<br>Vox. FPR: p = 0.13 | SI: **p = 0.005**<br>AVD: **p < 0.001**<br>Cl. F1: **p = 0.002**<br>Cl. TPR: **p < 0.001**<br>Vox. TPR: **p = 0.02**<br>Vox. FPR: **p = 0.04** |
| Case 2 |  | SI: p = 0.95<br>AVD: p = 0.16<br>Cl. F1: p = 0.33<br>Cl. TPR: p = 0.50<br>Vox. TPR: p = 0.09<br>Vox. FPR: **p = 0.02** | SI: p = 0.32<br>AVD: **p = 0.005**<br>Cl. F1: **p = 0.003**<br>Cl. TPR: p = 0.30<br>Vox. TPR: **p = 0.03**<br>Vox. FPR: p = 0.80 | SI: p = 0.53<br>AVD: p = 0.09<br>Cl. F1: p = 0.18<br>Cl. TPR: p = 0.30<br>Vox. TPR: p = 0.47<br>Vox. FPR: p = 0.89 | SI: p = 0.52<br>AVD: **p = 0.01**<br>Cl. F1: **p = 0.02**<br>Cl. TPR: p = 0.64<br>Vox. TPR: p = 0.06<br>Vox. FPR: p = 0.84 |
| Case 3 |  |  | SI: p = 0.25<br>AVD: p = 0.09<br>Cl. F1: **p = 0.03**<br>Cl. TPR: p = 0.89<br>Vox. TPR: p = 0.57<br>Vox. FPR: **p = 0.008** | SI: p = 0.38<br>AVD: p = 0.57<br>Cl. F1: p = 0.80<br>Cl. TPR: p = 0.15<br>Vox. TPR: p = 0.76<br>Vox. FPR: **p = 0.002** | SI: p = 0.09<br>AVD: **p = 0.01**<br>Cl. F1: **p = 0.01**<br>Cl. TPR: p = 0.97<br>Vox. TPR: **p = 0.03**<br>Vox. FPR: **p = 0.03** |
| Case 4 |  |  |  | SI: **p = 0.04**<br>AVD: **p = 0.004**<br>Cl. F1: **p = 0.007**<br>Cl. TPR: **p = 0.03**<br>Vox. TPR: p = 0.13<br>Vox. FPR: p = 0.16 | SI: p = 0.50<br>AVD: p = 0.25<br>Cl. F1: p = 0.64<br>Cl. TPR: p = 0.98<br>Vox. TPR: p = 0.38<br>Vox. FPR: p = 0.60 |
| Case 5 |  |  |  |  | SI: p = 0.05<br>AVD: **p = 0.04**<br>Cl. F1: p = 0.08<br>Cl. TPR: p = 0.08<br>Vox. TPR: p = 0.11<br>Vox. FPR: p = 0.96 |

Case 1: source-trained model applied on the target test dataset; Case 2: transfer learning; Case 3 : domain adversarial training of neural networks (DANN); Case 4: Semi-supervised DANN; Case 5: domain unlearning; Target-trained: target-trained model applied on the target test dataset.

### 3.1. Case 1: Train on the source domain and apply directly to the target domain

As shown in figure 4a, while the source-trained model detected most of the periventricular WMHs (PWMHs), it undersegmented their boundaries, and also missed some deep WMHs (DWMHs). From the boxplots in figure 5a and the performance metrics reported in table 1, case 1 showed the worst performance among all the cases. The segmentation was more precise in the low lesion load subjects (figure S3 in supplementary material) rather than high lesion load ones. This case showed significantly higher AVD and significantly lower cluster-wise F1-measure when compared to adversarial adaptation methods (cases 3-5) and showed significantly lower voxel- and cluster-wise TPR values when compared especially to DANN cases (cases 3 and 4) (table 2).

### 3.2. Case 2: Transfer the model trained on the source domain to the target domain with fine-tuning

The segmentation results improved with increased amounts of training data (figure 4). For instance, segmentation results with fine-tuning using 2 training subjects showed the worst results for both lesion loads, and improved with 12 and 18 training subjects. This is also evident from the heatmaps of the performance metrics shown for different numbers of training subjects and numbers of fine-tuned layers in figure 6. All the performance metrics showed best values for the training size of 18 subjects. The performance metrics were generally slightly lower when fewer layers in the decoder arm were fine-tuned, and also when there was less training data. However, the performance really started to increase when fine-tuning was done in the intermediate coarser layers (layers 3 and 4), and was noticeably higher when encoder layer 4 was fine-tuned, with even a small amount of training data. This is due to the rich domain-specific information at the intermediate layers. Therefore the visual results were better for the middle two columns of figure 4b when compared to their corresponding results when fine-tuned from layer 1 (the first column). The best performance metrics were obtained when the pretrained model is fine-tuned starting from layer 4 with 18 training subjects (table 1 and figure 5). In this case, TL provided better performance than case 1 and provided the highest median SI value among all cases. However, this case gave lower cluster-wise TPR values and higher AVD values when compared to other DA models (with the differences significant when compared to semi-DANN and target-trained cases, as reported in table 2).

**Figure 6:**
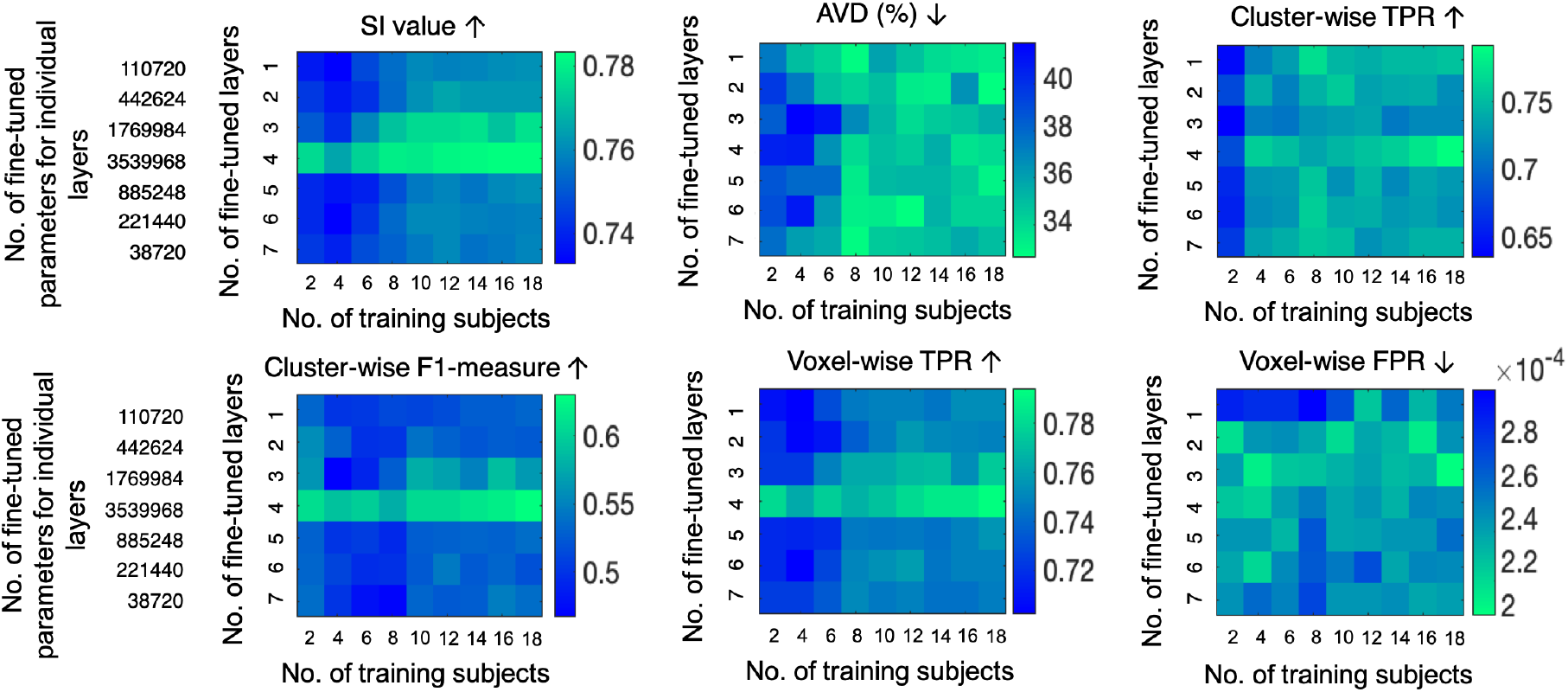
Heatmaps of performance metrics for the TL case (case 2) on the target test dataset, corresponding to the number of training subjects and the number of fine-tuned layers. The maps are shown for (top row, left to right) SI values, AVD (%), cluster-wise F1-measure, (bottom row, left to right) cluster-wise TPR, voxel-wise TPR and voxel-wise FPR values. The green end represents the best performance for all cases, ↑ shows that higher values indicate better performance and ↓ shows vice versa. Note that given a number of fine-tuned layers, the layers prior to them in the encoder end were frozen, and the remaining layers towards the decoder end were fine-tuned. The number of parameters associated with individual layers has been reported (only for a single plane). For example, if the final 5 layers are fine-tuned, the sum of the top 4 values in the left column denotes the total number of parameters fine-tuned per planar U-Net.

### 3.3. Case 3: Unsupervised domain adversarial training on source and target domains

In the case of unsupervised DANN, the segmentation was better than both cases 1 and 2, as shown in figure 4c on the target test dataset. The DANN model detected more lesion voxels along the ventricles and lesion edges when compared to the source trained model, and less false positives when compared to the TL case (especially in the low lesion load subjects). Even without using target labels for training, the model provided better delineation of PWMHs on the target test dataset, indicating the ability of the model to learn domain-invariant features. Unsupervised DANN gave the lowest voxel-wise FPR (significantly lower than semi-DANN), highest cluster-wise TPR and also the best voxel-wise TPR, on par with semi-DANN and target-trained cases. On the source test dataset, the DANN model achieved median SI = 0.91, AVD = 11.2%, cluster-wise F1-measure = 0.82, cluster-wise TPR = 0.90, voxel-wise TPR = 0.89 and voxel-wise FPR = 0.9 × 10^−4^. Also, DANN shows higher overlap between source and target feature representations at the layer before the label predictor, compared to the DU case, as shown in the t-SNE plot in figure 7b.

**Figure 7:**
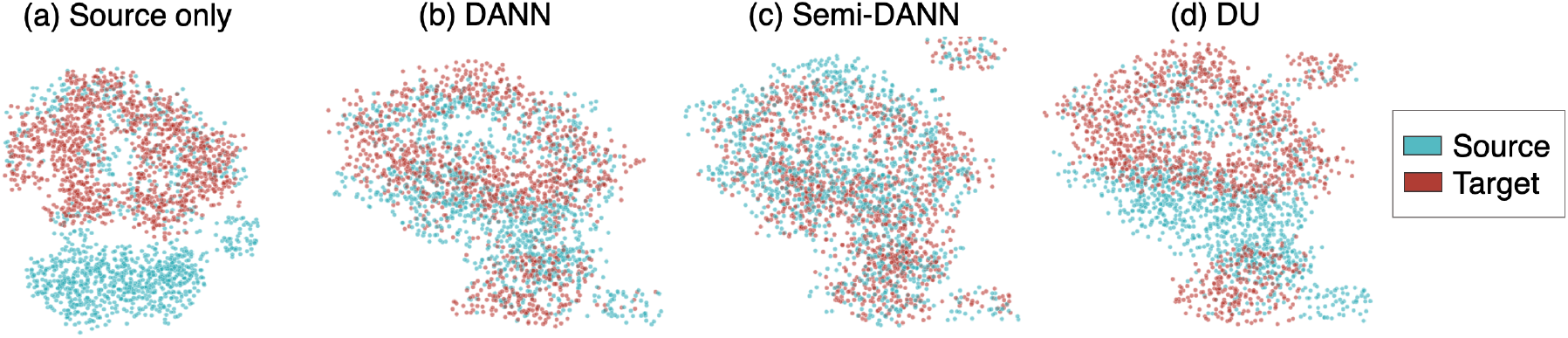
The effect of domain adaptation on the extracted feature distributions for source (blue) and target domains (red). T-distributed Stochastic Neighbour Embedding (T-SNE) plots of the feature map values at the layer before the label predictor for (a) model trained on source dataset only, (b) unsupervised DANN, (c) semi-DANN and (d) DU.

### 3.4. Case 4: Semi-supervised domain adversarial training (semi-DANN) on the source and target domains

In the case of semi-DANN, the addition of a fraction of the labelled target data to label prediction provided improvement in the segmentation performance over source-trained, TL and unsupervised DANN (figure 5 and table 1). Semi-DANN provided significantly higher cluster-wise F1-measure when compared to DANN, however with with significantly higher voxel-wise FPR as well. The improvement was subtle on visual assessment, and observable mainly along the boundaries of the PWMHs, as shown in figure 4d. The performance metrics achieved with semi-DANN method were not significantly different from those of target-trained model. On the source test dataset, the semi-DANN model achieved median SI = 0.86, AVD = 8.5%, cluster-wise F1-measure = 0.81, cluster-wise TPR = 0.87, voxel-wise TPR = 0.87 and voxel-wise FPR = 1.8 × 10^−4^. We observed that the semi-DANN also brings the distribution of extracted features from the source and target domains together with greater overlap, when compared to the unsupervised DANN, as shown in figure 7c.

### 3.5. Case 5: Iterative domain unlearning (DU) to remove scanner-bias between source and target domains

The DU model provided better performance than the source-trained case, but showed lower performance metrics compared to unsupervised DANN. However, none of the metrics were significantly different from case 3, except for higher voxel-wise FPR. On visual assessment, this case slightly oversegmented the PWMHs (figure 4e), mainly along the ventricles, which is also evident from the higher voxel-wise FPR values when compared to the unsupervised DANN. On the source domain test dataset, the DU model achieved SI = 0.83, AVD = 15.3%, cluster-wise F1-measure = 0.79, cluster-wise TPR = 0.84, voxel-wise TPR = 0.86 and voxel-wise FPR = 1.9 × 10^−4^. From the visualisation of feature representations in figure 7d, we can see that, for the DU model, while source and target features align as in the other DA cases, they still show domain gaps with slightly less mixing of features from different domains.

### 3.6. Train on the target domain from scratch and apply to the target domain

Out of all the test cases, the results from the model trained on the target training dataset showed the best segmentation performance on the target test dataset, as shown in figure 4f. But even in this case, the results showed a few false positive voxels in the high lesion load case, while the delineation of PWMHs was better than all the other cases. The target-trained model achieved the best cluster-wise F1-measure, cluster- and voxel-wise TPR values with the lowest AVD value as reported in table 1 and shown in figure 5, and significantly higher cluster-wise F1-measure and significantly lower AVD values when compared to the source-trained and TL cases. Interestingly, the source-trained model fine-tuned on the target dataset provided better median SI value (although with wider interquartile range in SI values) than the model trained from scratch on the target training dataset using same number of subjects. However, the other voxel- and cluster-wise metrics were better in target-trained case. Comparing with the adversarial training cases, unsupervised DANN (case 3) provided lower voxel-wise FPR values with on par cluster-wise TPR values (table 1).

## 4. Discussion and conclusions

In this work, we explored various domain adaptation techniques such as transfer learning, domain adversarial training and iterative domain unlearning for WMH segmentation using a triplanar ensemble model as the baseline method. In the case of TL, we also explored what would be the minimum number of subjects required for fine-tuning and which would be the best layers to fine-tune. We observed that domain adversarial training shows potential for better adaptation of the WMH segmentation task compared to other techniques on the given source and target datasets.

The source-trained model applied directly to the target test dataset achieved the worst performance out of all cases due to the differences in image resolution, pathology, intensity characteristics and lesion distribution, as shown in figure 3 (also refer to section 1 in supplementary material).

The performance of the TL case is better than training the baseline model on the combination of source and target dataset (refer to section 4 in supplementary material) on the target test dataset (even though the latter case shows better performance than case 1). For the TL case, when we varied the number of layers to fine-tune, the performance improved when more intermediate features from both encoders and decoders (layers 3 and 4) were fine-tuned (heatmaps in figure 6). Generally, initial convolutional layers often contain low-level features (e.g. edges) that tend to be naturally domain invariant, and hence fine-tuning this layer does not improve the target performance much, as shown in (Ghafoorian et al., 2017b). On the other hand, the intermediate layers (Zeiler and Fergus, 2014; Girshick et al., 2014) and the layers with coarsest features at the encoder end (Ghafoorian et al., 2017b) contain higher-level information such as lesion pattern that are domain specific. Hence, fine-tuning the coarsest layers (e.g. layer 4) provided better performance on the target test dataset. Also, it has been shown that fine-tuning initial encoder layers adds more training parameters and requires a larger number of training samples (50-100 subjects) to avoid over-fitting (unavoidable even with 25 subjects) (Ghafoorian et al., 2017b). Given the encoder-decoder architecture of U-Net, while fine-tuning more decoder layers led to the steady improvement in the performance, fine-tuning the initial layers of encoder reduced the performance, possibly due to the shortage of representative training data required for training the initial layers, as observed in Ghafoorian et al. (2017b). Regarding the number of subjects, we observed from the heatmaps that the performance started to improve when *>* 14 subjects were used for fine-tuning (starting from the coarser layer in the encoder). Using 18 subjects (the entire target training data) for fine-tuning greatly improved the results and provided performance metrics not significantly different from the unsupervised DANN case (except voxel-wise FPR). Although adding more representative training subjects (also depending on the variation in features between domains) could slightly improve the performance, we observed that in the case of smaller datasets, a minimum of 14-16 subjects for fine-tuning might provide good results.

We observed that the unsupervised DANN performed better than TL and the source-trained model. From figure 4b, c, d (also refer to supplementary figure S3) it can be seen that, even with a higher number of training subjects for TL, the DANN model extracted domain invariant features (e.g. contrast between lesion and background, distribution of PWMHs) and provided better visual results with less noise and more precise segmentation of boundary voxels. This was also reflected in the performance metrics, especially the cluster-wise metrics and AVD value, in table 1 and figure 5.

From figure 5 and table 1 and 2, we observed that the semi-DANN provided improvement over the unsupervised DANN on the target test dataset. Even though DANN achieved significantly lower cluster-wise F1-measure compared to the semi-DANN model, it also gave significantly lower FPR. Additionally, unsupervised DANN also provided voxel-wise TPR values on par with the semi-DANN, indicating that unsupervised DANN detects less false positives when compared to the semi-supervised case. Hence, while adding labelled target data might improve the performance of WMH segmentation in the target domain, it is necessary to weigh carefully the trade-off between improvement in the segmentation performance and the amount of manual effort involved, while choosing between unsupervised DANN and semi-DANN.

The DU model performed better than the source-trained model and provided performance metrics that were not significantly difference from the TL case. While training the DU model, we observed that the factor for weighting the domain confusion, *β*, plays a crucial role in achieving the domain invariance of the model. For the lower values of *β*, we found that the domain accuracy values were higher than 60% (where domain accuracy values closer to 50% are desirable indicating the maximal confusion of domains). We obtained the best results for the *β* value of 50 on the target dataset achieving a domain accuracy of 58%. Among the unsupervised models, DANN provided better performance than the DU model, with significant differences in voxel-wise FPR values. Also, DANN provided the better domain confusion with the domain accuracy of 47% at the layer before domain predictor (and a domain accuracy comparable to the DU model at the layer before the lesion label predictor as shown in the supplementary section 5). On visual assessment, the DU case oversegmented the PWMHs, while missing some DWMHs on the target test dataset, resulting in a lower cluster-wise TPR value.

From the t-SNE plots, it can be seen that with the model trained with the source data only, the source and target features are separable, while after domain adaptation the target features are aligned closer to the source features with a good overlap. The semi-DANN showed the maximum overlap of source and target feature representations at the layer before label predictor. The better overlap of domains with semi-DANN (compared to the unsupervised case) is expected due to the introduction of the labelled target data for training the label predictor in the semi-DANN. Among the unsupervised cases, DANN showed better overlap of the features when compared to the DU model. It is worth noting the performance of the adversarial training techniques (such as DANN and DU) depends mainly on the variations in lesion distribution and the acquisition characteristics between the source and target datasets (since they do not require lesion labels from the target dataset), rather than uncertainties in the manual segmentations on the target dataset (as in the TL case).

The size of source and target domain dataset is an important factor that affects the performance of domain adaptation techniques. The model transferred from source to target domain in case 2, uses both source and target domain datasets for pre-training and fine-tuning respectively and hence learns the characteristics from both datasets. On the other hand, the difference in the source and target domain datasets’ sizes and the inhomogeneity in inter-/intra-domain characteristics especially affect unsupervised methods like DANN. To better investigate this aspect, we chose the two best performing adversarial adaptation techniques from our test cases, DANN and semi-DANN, and trained them after swapping the datasets used for source and target domains. We observed that while DANN model is susceptible to slight changes in the performance (however, none of them significant except voxel-wise FPR), semi-DANN provided a consistent performance after domain swapping, without any significant difference in performance. For more details on the experiments and results, refer to supplementary section 3.

Concluding, we explored various DA techniques such as transfer learning and domain adversarial training techniques including DANN and DU. For the TL case, fine-tuning the intermediate layers towards the end of the encoder provided better performance than fine-tuning the initial layers. The DANN models performed better than TL and the DU model on our target dataset. Particularly, the semi-DANN provided the best performance with significant improvements over DU and TL cases. However, even without the addition of labelled target training data, the unsupervised DANN provided better performance compared to TL and DU, and results on par with the semi-DANN.

## Acknowledgements

This research was funded in whole, or in part, by the Wellcome Trust [Grant number 203139/Z/16/Z]. For the purpose of open access, the author has applied a CC BY public copyright licence to any Author Accepted Manuscript version arising from this submission. This work was also supported by the Engineering and Physical Sciences Research Council (EPSRC) and Medical Research Council (MRC) [grant number EP/L016052/1] and Wellcome Centre for Integrative Neuroimaging, which has core funding from the Wellcome Trust. The computational aspects of this research were funded from National Institute for Health Research (NIHR) Oxford BRC with additional support from the Wellcome Trust Core Award Grant Number 203141/Z/16/Z. The Oxford Vascular Study is funded by the National Institute for Health Research (NIHR) Oxford Biomedical Research Centre (BRC), Wellcome Trust, Wolfson Foundation, the British Heart Foundation and the European Union’s Horizon 2020 programme (grant 666881, SVDs@target). The views expressed are those of the author(s) and not necessarily those of the NHS, the NIHR or the Department of Health. VS is supported by the Wellcome Centre for Integrative Neuroimaging. GZ is supported by the Italian Ministry of Education (MIUR) and by a grant “Dipartimenti di eccellenza 2018-2022”, MIUR, Italy, to the Department of Biomedical, Metabolic and Neural Sciences, University of Modena and Reggio Emilia. ND is supported by the Engineering and Physical Sciences Research Council (EPSRC), Medical Research Council (MRC) (EP/L016052/1). PMR is in receipt of a NIHR Senior Investigator award. LG is supported by the Monument Trust Discovery Award from Parkinsons UK (Oxford Parkinsons Disease Centre, J-1403), the MRC Dementias Platform UK (MR/L023784/2) and the National Institute for Health Research (NIHR) Oxford Health Biomedical Research Centre (BRC). MJ is supported by the NIHR Oxford Biomedical Research Centre (BRC).

We acknowledge all the participants. For the NDGEN dataset, we are grateful to Prof. Gordon K. Wilcock and all the staff of Oxford Project to Investigate Memory and Ageing (OPTIMA) study. For the OXVASC dataset, we acknowledge the use of the facilities of the Acute Vascular Imaging Centre, Oxford. We also thank Dr. Chiara Vincenzi and Dr. Francesco Carletti for their help on generating the manual masks used in our experiments.

MJ receives royalties from licensing of FSL to non-academic, commercial parties. The authors report no potential conflicts of interest.

## Footnotes

2 TrUE-Net code available in https://git.fmrib.ox.ac.uk/vaanathi/true_net_wmh_segmentation_pytorch

3 3DU code available at: https://github.com/nkdinsdale/Unlearning_for_MRI_harmonisation

